# Identification of novel inhibitors targeting serine acetyltransferase from *Neisseria gonorrhoeae*

**DOI:** 10.1101/2024.12.03.626642

**Authors:** Keely E.A. Oldham, Wanting Jiao, Erica Prentice, Joanna L. Hicks

## Abstract

*Neisseria gonorrhoeae* is an obligate human pathogen and the etiological agent of the sexually transmitted infection, gonorrhoea. The rapid emergence of extensively antimicrobial-resistant strains, including those resistant to all frontline antibiotics, has led to *N. gonorrhoeae* being labelled a priority pathogen by the World Health Organization, highlighting the need for new antimicrobial treatments. Given its absence in humans, targeting *de novo* cysteine biosynthesis has been identified as a promising avenue for developing new antimicrobials against drug-resistant bacteria. The biosynthesis of cysteine is catalyzed by two enzymes; serine acetyltransferase (SAT/CysE) which catalyzes the first step and *O*-acetylserine sulfhydrylase (OASS/CysK) that catalyzes the second step incorporating sulfur to form L-cysteine. CysE is reported to be essential for bacterial survival in several bacterial pathogens including *N. gonorrhoeae*. Here, we have conducted virtual inhibitor screening of commercially available compound libraries against SAT from *N. gonorrhoeae* (*Ng*SAT). We have identified a hit compound with an IC_50_ of 13.9 µM and analyzed its interactions with the enzyme’s active site. This provides a platform for the identification and development of novel SAT inhibitors to combat drug-resistant bacterial pathogens.

## Introduction

Antimicrobial resistance is an ever-growing global problem and has been declared a public health crisis. *Neisseria gonorrhoeae* (gonococcus) is a Gram-negative bacterium and an obligate human pathogen, causing the sexually transmitted infection (STI), gonorrhoea. Globally, there are approximately 87 million cases annually, making it the second most common bacterial STI, after chlamydia (1). Concerningly, *N. gonorrhoeae* can readily acquire antimicrobial-resistance, with documented resistance emerging for every class of antibiotic used for its treatment since the introduction of sulfonamides in the 1930s (2). In 2017, *N*. *gonorrhoeae* was labelled as a priority pathogen by the World Health Organization, (3) and in the absence of an effective vaccine (4), coupled with the increasing prevalence of antibiotic-resistant strains, there is a urgent need for new antimicrobial treatments.

An emerging theme in the development of new antimicrobials is the targeting of key metabolic pathways that are important for infection and virulence. As such, there is expanding interest in targeting amino acid biosynthetic pathways in bacteria. One such pathway is the *de novo* synthesis of the amino acid L-cysteine. This pathway is well-conserved across bacteria and plants but is notably absent in mammals (5). Alongside being synthesis of thiol-containing metabolites, such as glutathione and thioredoxin, both of which are important in regulating the cell redox state and for protection from oxidative stress encountered during infection (5). Cysteine is synthesized via a two-step reaction by the cysteine biosynthetic enzymes, serine *O*-acetyltransferase (SAT/CysE, EC 2.3.1.30) and *O*-acetylserine sulfhydrylase (OASS-A/OASS-B; CysK/CysM, EC 4.2.99.8) (6, 7). SAT catalyzes the acetylation of serine using acetyl-CoA to produce *O*-acetylserine which is subsequently condensed with sulfide by CysK, or thiosulfate in the case of CysM, to form L-cysteine (Fig 1).

**Fig 1.**
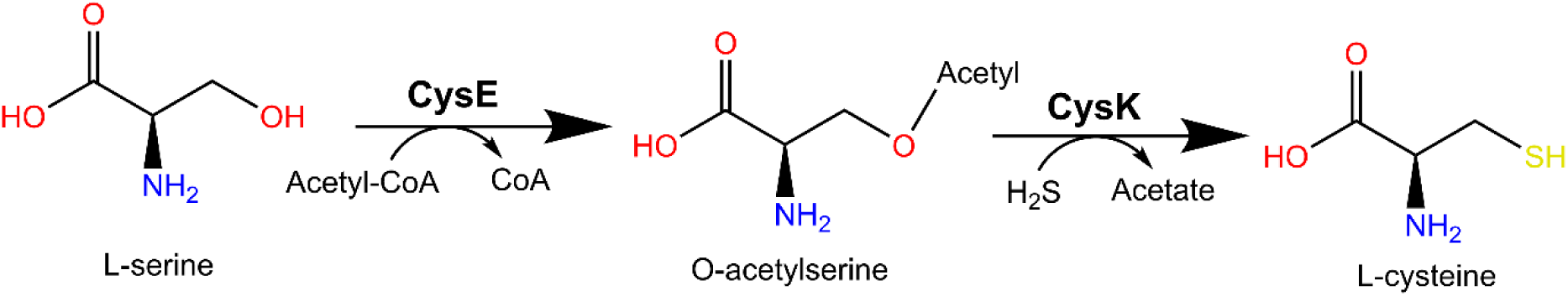
*De novo* cysteine biosynthesis reaction in *Neisseria gonorrhoeae.* CysE, serine *O*-acetyltransferase (SAT); CysK, *O*-acetylserine sulfhydrylase (OASS-A).

The *de novo* synthesis of cysteine is tightly regulated at the protein level which is primarily through feedback inhibition of SAT by cysteine, as high concentrations can lead to damaging Fenton chemistry (8). Additionally, cysteine synthesis regulates cellular sulfur flux through the formation of the cysteine synthase complex (CSC). The CSC is formed by the reversible association of one SAT hexamer and two CysK dimers formed through the SAT C-terminal tail binding into and occluding the CysK active site, enhancing SAT activity while abolishing CysK activity (7, 9). CSC formation is favored under low sulfide concentrations (9) which allows *O*-acetylserine to accumulate and dissociate the CSC (7).

The absence of the cysteine biosynthesis pathway in mammals makes it ideal for antimicrobial targeting by reducing the chance of off-target effects (10). Additionally, targeting of SAT has been reviewed (11) and reported for notable antimicrobial resistant bacterial pathogens including, *Escherichia coli* (12), *Salmonella enterica* Serovar typhimurium (13) *Staphylococcus aureus* (14) and *Mycobacterium tuberculosis* (15). SAT from *N. gonorrhoeae* (*Ng*SAT) has been shown to be differentially expressed during the stringent response (16) and is an essential gene (17). Thus, we propose *Ng*SAT is a promising new target for antimicrobial development. Previously we kinetically and structurally characterized *Ng*SAT (PDB 6WYE and 7RA4), demonstrating that it has comparable activity to other bacterial SAT homologs and possesses a conserved structural fold and active site residues (18). Here, we have used structure-based virtual inhibitor screening of commercial compound libraries against *Ng*SAT and tested *in vitro* 28 hit compounds with one compound (compound 2) identified as a potent inhibitor of *Ng*SAT.

## Results and Discussion

### Conservation of NgSAT (CysE) amongst *Neisseria gonorrhoeae* isolates

Conservation of active site residues is critical for ensuring broad spectrum activity against different *N. gonorrhoeae* isolates. Analysis of the presence and conservation of SAT (*cysE*) amongst *N. gonorrhoeae* strains was conducted via BLAST using *cysE* from *N. gonorrhoeae* MS11 as a query against all available *N. gonorrhoeae* genomes in the NCBI database. *cysE* is highly conserved in *N. gonorrhoeae* (99.88-100% sequence identity) over the length of the sequence (819 nucleotides, 272 residues) with the majority of strains having an identical isoform (156/157 genomes). To explore the distribution of SAT amongst *N. gonorrhoeae* isolates, we expanded our search to include all available *N*. *gonorrhoeae* genomes in the PubMLST database. A *cysE* gene (NEIS0501) was annotated in in all isolate genomes (19,729 genomes) and is well-conserved, with 97.7-100% similarity reported. Furthermore, comparison of these isolates shows that the majority genomes had an annotated *cysE* identical to the MS11 *cysE* query (18,978/19,729, 96% of genomes had an identical copy). Overall *cysE* is strictly conserved across *N. gonorrhoeae* reference strains and circulating isolates, thus making it a suitable target for antimicrobial applications.

### *Ng*SAT active site model with both substrates bound

The crystal structure of *Ng*SAT (PDB 6WYE) contains malate in the serine binding site and lacks acetyl-CoA. To examine substrate-enzyme interactions, we generated an active site model with both substrates bound. The crystal structures of *Yersinia pestis* SAT (PDB 3GVD) and *Haemophilus influenzae* SAT (PDB 1SST) contain the inhibitor cysteine and the product co-enzyme A in the active site, respectively. With the structural similarity between serine and cysteine these two structures were used as the starting point for positioning serine and acetyl-CoA into the active site of *Ng*SAT. The initial active site model of *Ng*SAT with both serine and acetyl-CoA placed was then further optimized through a series of enzyme minimization, refinement, and quantum mechanics/molecular mechanics (QM/MM) optimization.

The active site model (Fig 2A) shows that both substrates can bind simultaneously. The nucleophilic oxygen atom of serine is 3.9 Å away from the acetyl carbon of acetyl-CoA, positioning it for the reaction. Acetyl-CoA primarily interacts with the active site via polar interactions from its phosphate groups (Fig 2B). The phosphate forms salt bridge interactions with two Lys246 residues from the two chains forming the active site. The diphosphate group establishes a salt bridge interaction with Lys223. Additionally, acetyl-CoA forms hydrogen bond (H-bond) interactions with the backbones of residues Ala226, Ala208, and Ala249 (Fig 2B). The binding of serine is stabilized by interactions between its backbone and charged residues Arg196, Asp96, and Asp161 (Fig 2C). The side chain of serine interacts with two histidine residues, His197 and His162. The current active site model suggests that His162 will deprotonate serine for the reaction.

**Fig 2.**
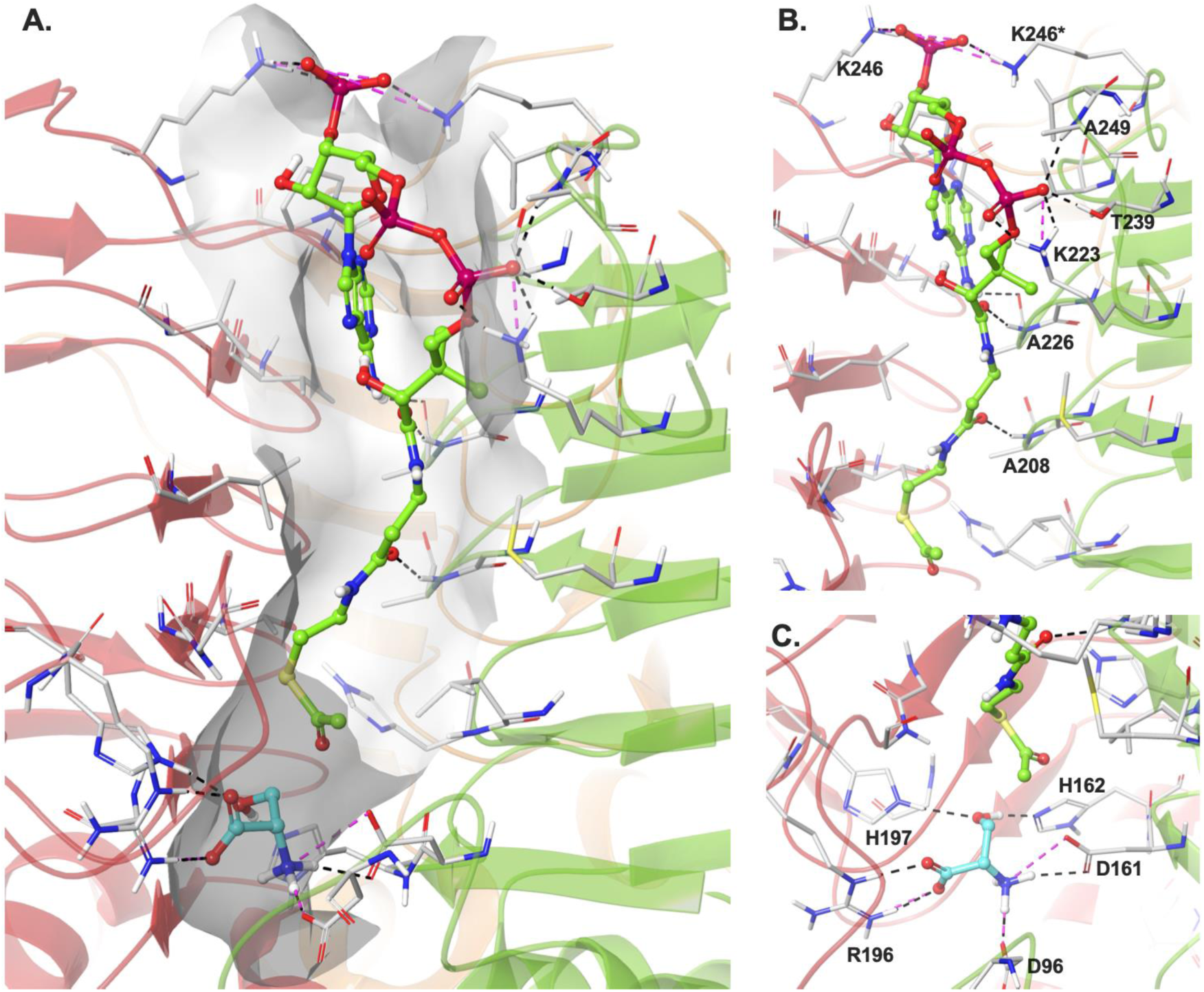
Active site model of *Ng*SAT with both serine and acetyl-CoA bound. (A) Binding sites and relative positions of serine (cyan carbon) and acetyl-CoA (green/pink carbon). (B) and (C) binding sites for acetyl-CoA and serine, respectively. Residues involved in polar interactions are labelled. H-bond interactions are displayed as black dashed lines and salt bridges are displayed as magenta dashed lines.

### Hit inhibitor screening and characterization

We employed a structure-based virtual screening approach to search for potential *Ng*SAT receptor for virtual screening. Compounds with drug-like properties were obtained from the ZINC database (19) which, after preparation, contained a total of ∼10.5 million compounds. The drug-like compound library was screened against the substrate-bound crystal structure model of *Ng*SAT, with the acetyl-CoA binding site being the primary target binding site for screening.

The top 1000 poses from the screening calculation were examined for enzyme-ligand interactions. Compounds showing interactions to the backbones of Ala208 and Ala226, and additional interactions to residues in the acetyl-CoA binding site were selected for further molecular mechanics with generalized Born and surface area calculations (MM-GBSA) (54 compounds). During the MM-GBSA calculations, active site residues within 5 Å of the docked compound were refined to optimize the enzyme-ligand interactions. An MM-GBSA binding free energy was also computed for each compound. Following the MM-GBSA calculation, compounds that had a predicted binding free energy below -50 kcal/mol and retained the interactions with the backbones of Ala208 and Ala226, were selected for validation using short (2 ns) molecular dynamics (MD) simulations (23 compounds). The MD trajectories of the ligand-enzyme complex were examined and compounds that maintained their interactions during the MD simulations were recommended for testing (nine compounds).

In addition to the drug-like compound library, a second library containing large compounds with a molecular weight greater than 500 Daltons was screened. The compounds were retrieved from the ZINC database (19) and prepared for docking. The prepared library contained ∼1.3 million compounds. The target receptor conformation used for screening the big-compound library was a representative active site conformation obtained from a 5 μs MD simulation of the *Ng*SAT enzyme. The acetyl-CoA site was the target screening site. The top 1000 compounds were examined using criteria similar to those mentioned above; 152 compounds underwent MM-GBSA calculation, of which 82 maintained backbone interactions with the alanine residues and were predicted to have binding free energies below -50 kcal/mol. The 82 compounds were clustered into 36 groups based on their structures, and one representative compound from each group was recommended for further testing.

Of the 46 total recommended compounds, 28 could be purchased at the time of the study. The details of hit compounds experimentally tested can be found in Supplementary Table 1.

### *In vitro* screening and characterization

Hit compounds identified from virtual inhibitor screening were experimentally evaluated by measuring inhibition of recombinant *Ng*SAT activity. Recombinant *Ng*SAT was expressed and purified following published methods (18). Previously, we characterized *Ng*SAT using a direct assay at 232 nm to determine the Michaelis-Menten kinetic parameters for substrates serine and acetyl-CoA (18). Given the high absorbance of aromatic compounds at this wavelength, *Ng*SAT inhibitor screening was performed using a plate-based coupled assay detecting production of CoA using 5,5’-Dithio-bis-(2-nitrobenzoic acid) (DTNB) through measuring production of 2-nitro-5-thiobenzoate (TNB) adapted from (13). *Ng*SAT activity was determined from a co-enzyme A DTNB standard curve (Supplementary Fig 1).

For *in vitro* screening, 28 of the 46 total compounds recommended were available and ordered from commercial vendors. Stocks of 1 mM were prepared for screening in 100% dimethyl sulfoxide (DMSO). Initial screening at 100 µM in the presence of 1 mM serine and 0.15 mM acetyl-CoA was conducted to identify inhibition and check solubility (Fig 3A).

**Fig 3.**
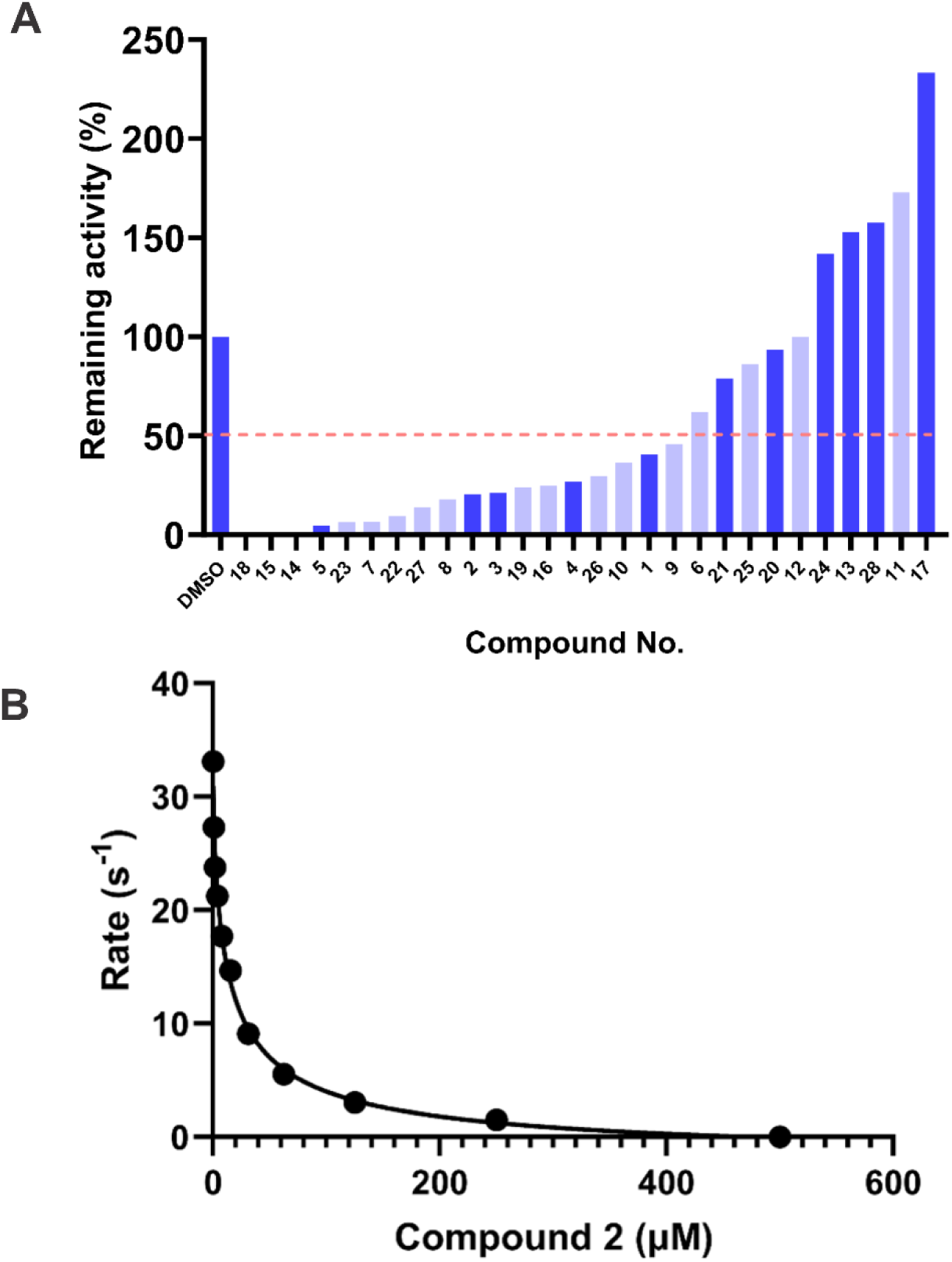
Screening of inhibitors against NgSAT (A)100 µM activity screen of library compounds against *Ng*SAT. Remaining activity (%) is calculated as rate with inhibitor present divided by rate in presence of DMSO (V_i_/V_o_) multiplied by 100. Soluble and insoluble compounds are highlighted in dark and light blue, respectively. Dashed pink line corresponds to 50% activity cut-off for downstream testing. Compounds 14, 15 and 18 have negative rates and for display purposes are represented as zero. Well containing DMSO only treated as positive control (100% activity). (B) IC_50_ dose response curve for Compound 2. *Ng*SAT activity measured in presence of 0.1 mM acetyl-CoA and 0.5 mM serine. Rate (s^-1^) was calculated as per hexamer. Datapoints are single replicates R^2^ =0.9924

Compounds that showed more than 50% inhibition and did not precipitate in the assay reaction were subjected to IC_50_ screening (compounds 1, 2, 3, 4 and 5). Upon further testing, compounds 1, 3, 4 and 5 did not display dose-dependent inhibition and therefore were classified as false positives from the original screen (data not shown). Compound 2, however, demonstrated inhibition (20.6% remaining activity at 100 µM) and collection of a dose response curve produced an IC_50_ of 13.9 ± 4.3 µM (R^2^ =0.9924) (Fig 3B), making it a potent hit compound.

### Docking Interactions for Compound 2

Compound 2 (1-(6-ethoxy-4-methyl-2-quinazolinyl)-N-[2-(1H-imidazol-1-yl)ethyl]-3-piperidinecarboxamide), consists of; an imidazole ring, with an amide linker connected to a piperidine ring, which itself is connected to a quinazoline ring with an ether attached (Figure 4) The compound does not violate any Lipinski’s rules (Table 1) and has qualities consistent with a drug-like compound (20).

**Figure 4.**
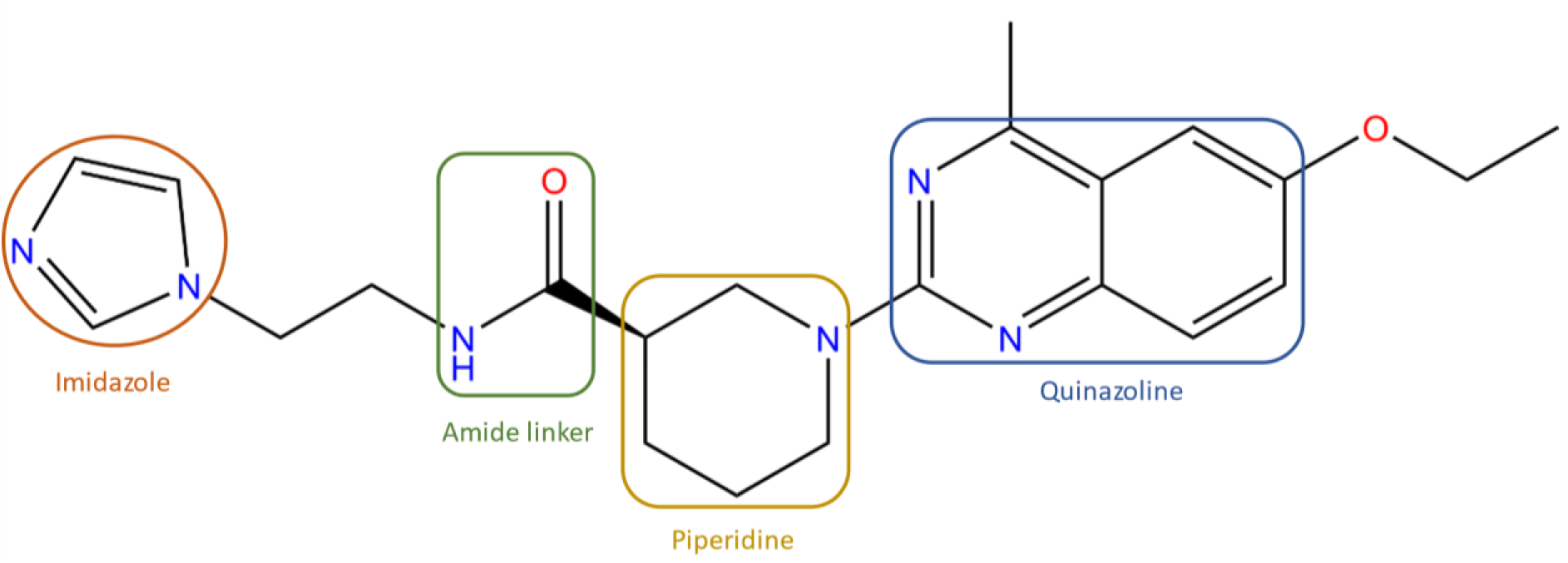
Structure of Compound 2 (1-(6-ethoxy-4-methyl-2-quinazolinyl)-N-[2-(1H-imidazol-1-yl)ethyl]-3-piperidinecarboxamide). Functional groups imidazole, piperidine and quinazoline are highlighted in orange, yellow, and blue, respectively. Figure created using ChemDraw Prime.

**Table 1.** Compound 2 chemical properties. tPSA, polar surface area (Å^2^); logP, partition coefficient.

| Chemical Formula | Net charge | H-bond donors | H-bond acceptors | Molecular Weight (g/mol) | logP | tPSA | Rotatable bonds | Apolar desolvation | Polar desolvation |
| --- | --- | --- | --- | --- | --- | --- | --- | --- | --- |
| C<sub>22</sub>H<sub>28</sub>N<sub>6</sub>O<sub>2</sub> | 0 | 1 | 7 | 408.5 | 2.566 | 85 | 7 | 9.87 | -17.44 |

Examination of *Ng*SAT poses with compound 2 docked from virtual inhibitor screening, show compound 2 forms notable active site interactions with both the serine and acetyl-CoA binding sites (Fig 5). There are four H-bond interactions formed between adjacent monomers with serine binding site residue Asp161 (Chain C) and with backbone interactions with acetyl-CoA binding residues Ala208 (Chain C), Ala226 (Chain C), and Gly188 (Chain A) (Fig 5B), and is complemented with a number of hydrophobic interactions from the acetyl-CoA site with the piperidine and quinazoline ring (Fig 5C). Additionally, these residues that interact with serine and acetyl-CoA site are strictly conserved, not only across *Neisseria* but also Proteobacteria (Fig 6). Furthermore, compound 2 does not share a high degree of similarity with published SAT inhibitors based on Tanimoto similarity scores (Fig 7) (scores lower than 0.7 (21)), making compound 2 a novel SAT inhibitor.

**Fig 5.**
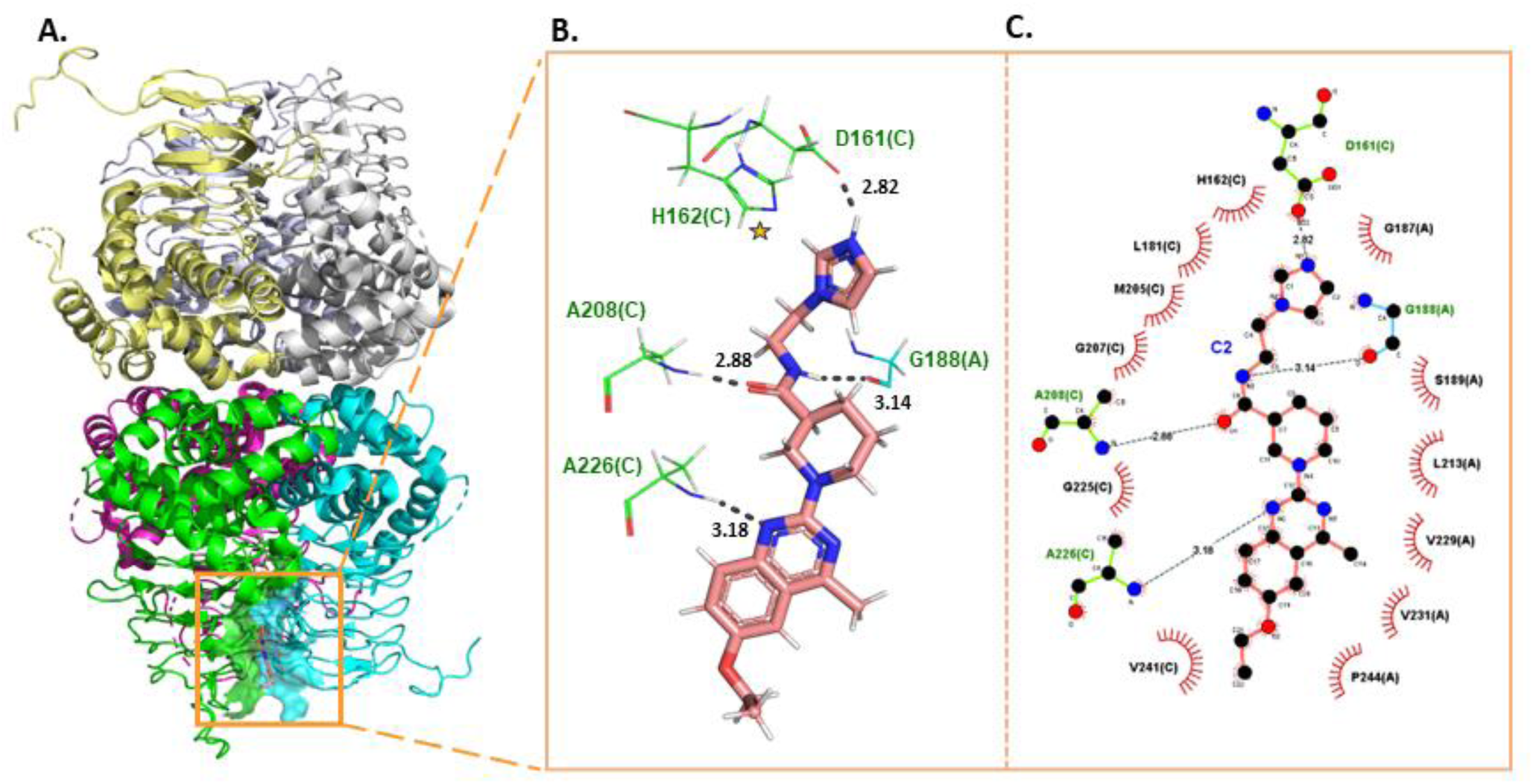
Docked interactions of compound 2 binding to *Ng*SAT. (A) *Ng*SAT hexamer (6WYE) with compound 2 forms hydrogen bond interactions with *Ng*SAT (PDB: 6WYE) serine and acetyl-CoA binding pockets (Chain A and Chain C, colored cyan and green, respectively). (B) Hydrogen bond interactions (gray) between compound 2 (pink sticks) and *Ng*SAT residues (green and cyan wires). (C) Two-dimensional LigPlot schematic of H-bonds and hydrophobic interactions between compound 2 and *Ng*SAT residues (bond lengths not to scale). H-bonds are shown as black dashes, and hydrophobic interactions are represented as red radiating lines. Catalytic histidine 162 (H162) is indicated stars. All bond lengths are reported in Angstroms (Å). Figures created using PyMOL and LigPlot^+^ (22).

**Fig 6.**
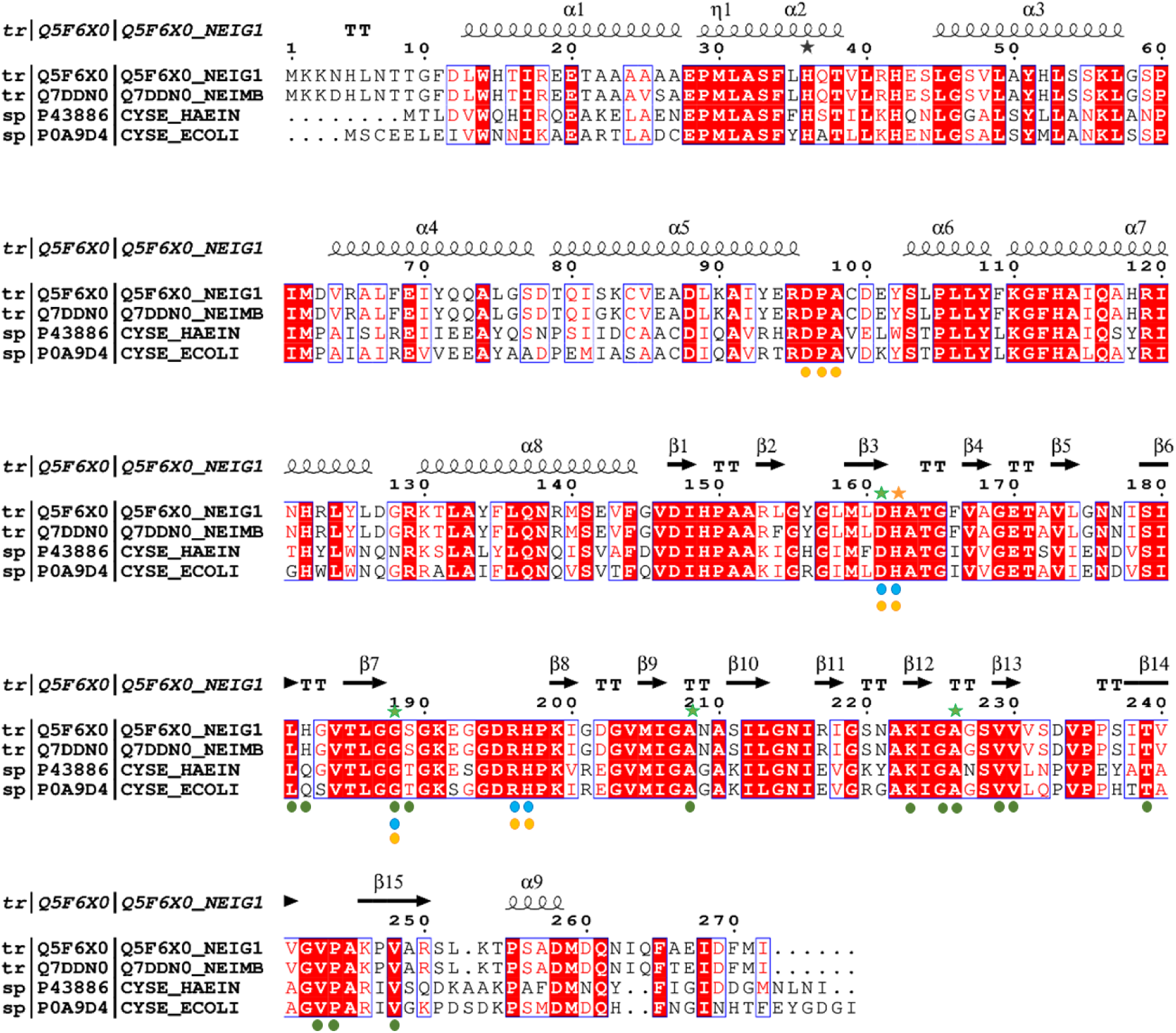
Multiple sequence alignment of bacterial SAT isoforms. Serine, acetyl-CoA and cysteine binding residues are annotated as green, blue and orange dots, respectively. Compound 2 hydrogen bond interactions are annotated by green stars. Active site catalytic His162 is highlighted as orange star. Sequences: NEIG1, *Neisseria gonorrhoeae* (PDB 6WYE); NEIMB, *Neisseria meningitidis* (no crystal structure); HAEIN, *Haemophilus influenzae* (PDB 1SST); ECOLI, *Escherichia coli* (PDB 1T3D).

**Fig 7.**
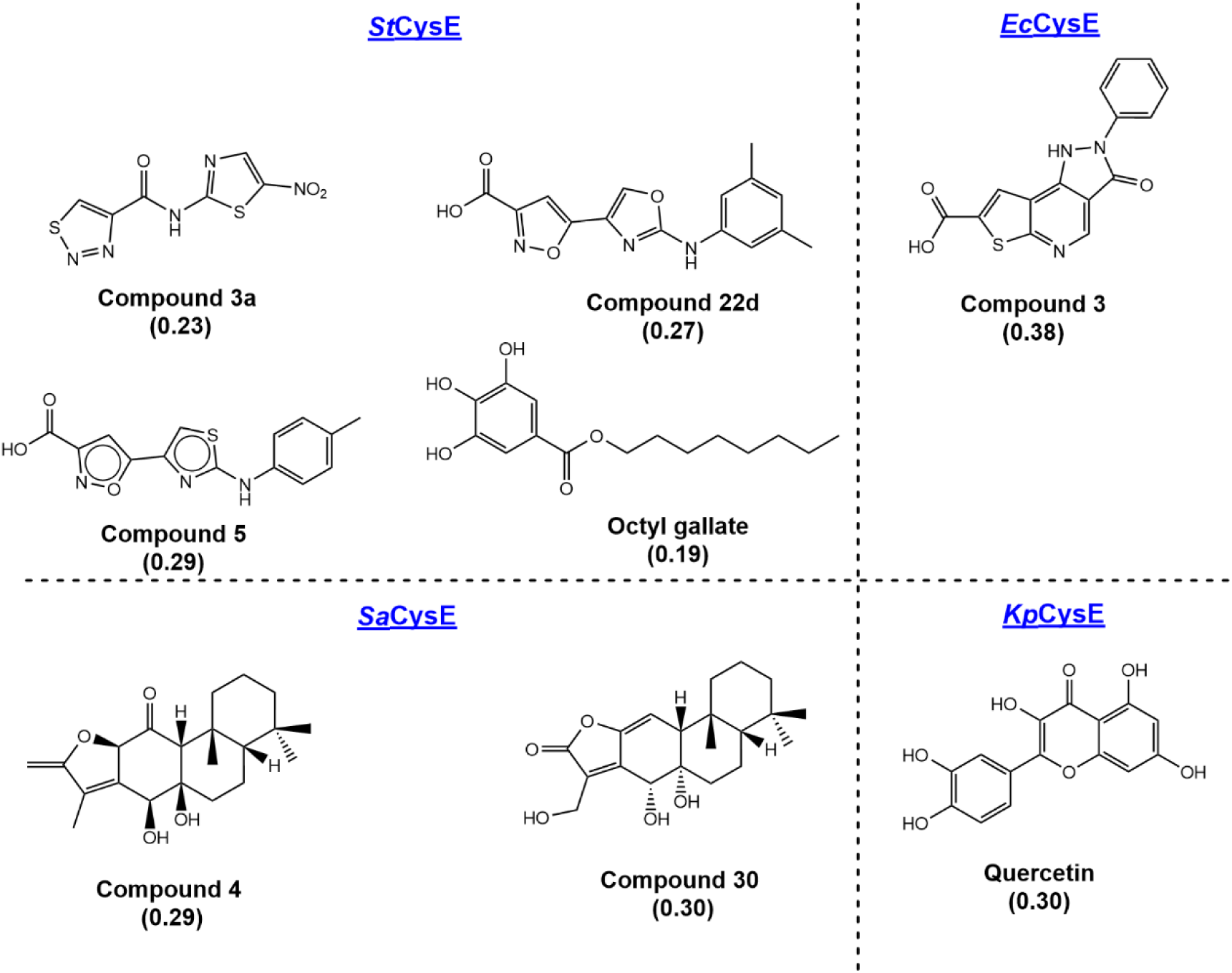
Characterized inhibitors against bacterial SAT homologs. Tanimoto fingerprints co-efficient similarity scores of reported SAT inhibitors compared to compound 2 are shown in brackets. Tanimoto scores were calculated using RDKit (23).

## Discussion

Antimicrobial-resistant *N. gonorrhoeae* is a global burden on human health, and in the absence of an effective vaccine, antimicrobials remain the front-line tool for managing gonococcal disease. The targeting of the *de novo* cysteine biosynthesis pathway has been highlighted as an avenue for development of antimicrobial compounds and adjuvants given its role in mitigation of oxidative stress and the absence of the pathway in humans. Here we report the first inhibitor of *Ng*SAT and demonstrate that structure-based virtual inhibitor screening is a successful strategy for identifying inhibitors. After computationally screening two libraries of 1.3 and 10.5 million compounds respectively from the ZINC15 database, 46 compounds were identified that met our criteria *in silico*, and of the 28 compounds tested experimentally, the best compound to inhibit *Ng*SAT activity *in vitro* was compound 2. Docking analysis revealed notable H-bond and hydrophobic interactions between compound 2 and both *Ng*SAT serine and acetyl-CoA binding pockets along the length of the compound. Dose-response analysis of compound 2 showed it to be a promising inhibitor with an IC_50_ of 13 µM.

The use of both virtual and experimental testing is needed for the rapid identification of novel inhibitors and has proved to be a successful strategy for identifying bacterial SAT inhibitors. The inhibition observed for compound 2 is in the low micromolar range, making it one of the better SAT inhibitors reported to date (11). Although inhibitors that exhibit sub-micromolar inhibition have been reported, ensuring that the compound is membrane permeable remains a challenge, particularly given the cytoplasmic nature of SAT isoforms, including *Ng*SAT, (24) and is a notorious problem for Gram-negative bacteria (25). Therefore, determining the MIC of compound 2 against *N. gonorrhoeae* is needed to ensure the compound is bioactive and thus a viable antimicrobial.

As highlighted previously (11), *in vivo* validation of SAT inhibitors has often been limited in their ability to disrupt bacterial growth. Only recently, Toyomoto and colleagues showed SAT inhibition in *E. coli* could not only disrupt bacterial growth, but also deplete intracellular concentrations of thiol-metabolites, as well as increase sensitivity to antibiotics and oxidative stress (12). This highlights the multi-faceted role SAT serves within bacteria, indicating that not only will *de novo* cysteine synthesis be disrupted by inhibition, but also resistance to oxidative stress, and existing resistance to known antibiotics.

A caveat of targeting of *de novo* cysteine synthesis is the possibility of resistance via up-regulation of alternative sulfur acquisition and assimilation pathways to obtain cysteine (12). Sulfur acquisition in *N. gonorrhoeae* is unique in that it possesses a sulfate/thiosulfate ABC transporter (26), thiol amino acid transporters (methionine (27), cysteine, and cystine (27, 28)), but lacks importers for thiol compounds taurine or glutathione (29). Additionally, glutathione is maintained at comparatively high levels in *N. gonorrhoeae* (15 mM vs. 5 mM in *E. coli*) (30), and given the absence of glutathione importers or the glutathione recycler γ-glutamyl transpeptidase (31), *N. gonorrhoeae* is reliant on *de novo* synthesis of glutathione and would rely on cysteine to maintain these high glutathione concentrations. Given this lack of pathway redundancy and high need for antioxidant defense during infection, and the essential nature of *cysE*, *N. gonorrhoeae* is less likely to develop resistance to a SAT inhibitor making it an ideal organism for *de novo* cysteine inhibition.

In summary, we have demonstrated that structure-based virtual inhibitor screening is a promising avenue for identifying inhibitors of *Ng*SAT. We report compound 2 to show inhibition of *Ng*SAT in the low micromolar range making it one of the most potent SAT inhibitors to date. Furthermore, given the low similarity to published bacterial SAT inhibitors compound 2 is a novel SAT inhibitor. Given the essential nature of *cysE* in *N. gonorrhoeae*, compound 2 is a promising candidate for antimicrobial development against this pathogen.

## Materials and Methods

### Bioinformatic analysis of *Ng*SAT

Investigation of the conservation and distribution of SAT sequences in *N. gonorrhoeae* was conducted through a BLASTN search of the complete *N. gonorrhoeae* genomes (taxid: 485) held by NCBI (“Complete prokaryote Genome Database”) using *Ng*SAT nucleotide sequence from *N. gonorrhoeae* strain MS11 as a query (RefSeq Accession; WP_025457874.1, locus_tag NGFG_RS07905, NC_022240.1:1458601-1459419). Determining SAT presence in *N. gonorrhoeae* isolates (24,173 isolates) held in the PubMLST (accessed Jan 2024) (32) was conducted via searching the SAT sequence from *N. gonorrhoeae* strain MS11 as a query (RefSeq Accession; WP_003989278.1, (NC_022240.1:1458601-1459419), NEIS0501) using the BLASTN search plug-in.

### Substrate-bound structural model of *Ng*SAT

A hexamer of *Ng*SAT was generated from symmetry operations on the crystal structure of *Ng*SAT (PDB 6WYE). The malate molecules were deleted. Crystal structures of *Yersinia pestis* SAT (PDB 3GVD), which contains cysteine in the serine site, and *Haemophilus influenzae* SAT (PDB 1SST), which contains co-enzyme A, were aligned with the *Ng*SAT crystal structure at chains A and C (*Ng*SAT chain names). The cysteine and co-enzyme A molecules binding in the A/C interface were extracted from 3GVD and 1SST and merged with the *Ng*SAT crystal structure. The cysteine ligand in the active site was then mutated to serine *in silico* in Maestro from Schrödinger Suite (33). Thus, a structure of *Ng*SAT with CoA and serine bound in the active site was generated. This merged structure was then prepared for modelling using the protein preparation wizard in the Schrödinger Suite (34, 35). During protein preparation, hydrogen atoms were added, ionization states for the ligands were generated using Epik (36-38) for pH 7±2, and all water molecules were deleted. Hydrogen bond pairs were optimized, and protonation states of ionizable residues were assigned using PROPKA (34, 39) for pH 8.0. The structure was then subjected to energy minimization, converging heavy atoms to an RMSD of 0.3 Å. An acetyl group was then built onto the CoA molecule, followed by another round of restrained minimization, converging heavy atoms to an RMSD of 0.3 Å.

The active site model of *Ng*SAT containing serine and acetyl-CoA in the active site in the A/C interface was further optimized using Prime refinement (40-42). Atoms within 8 Å of the serine and acetyl-CoA substrates were minimized to sample conformational changes around the serine and acetyl-CoA binding site to better accommodate the ligand molecules that were absent in the initial crystal structure. Finally, QM/MM optimization calculation was set up using QSite (43:Philipp, 1999 #449) from the Schrödinger Suite (44), to search for catalytic-relevant conformations (positionings of the serine and acetyl group in relation to the catalytic dyad). The QM region was defined to include side chains of His162, Asp161, serine substrate, and part of the acetyl-CoA molecule defined by inserting a hydrogen cap between C6P and C7P on acetyl-CoA. The resultant QM region contained 51 atoms. The QM optimization was set up using the DFT-B3LYP method and lacvp* basis set. The MM region was minimized using the Truncated Newton algorithm with a maximum of 1000 cycles, and the QM region was minimized with a maximum of 5000 iterations.

### Molecular dynamics simulation of *Ng*SAT

The crystal structure of *Ng*SAT contains missing atoms in disordered regions. Before MD simulations could be set up, a complete structure containing residues 1 to 264 was generated by building in the missing residues and side chains in COOT(45), fitting to the electron density map. The complete hexamer was then generated based on crystal symmetry. The malate molecules were deleted. Crystal structures of *Yersinia pestis* SAT (PDB 3GVD) and *Haemophilus influenzae* SAT (PDB 1SST) were aligned to the top and bottom trimers of *Ng*SAT (as there appears to be a degree of rotation relative to the top and bottom trimers in *Ng*SAT compared to the other two crystal structures). Then, all six cysteine and the four CoA molecules in the “open” active sites of *Ng*SAT were extracted from 3GVD and 1SST and merged with the *Ng*SAT structure. The cysteine ligands in the active sites were then mutated to serine *in silico*. Thus, a structure of the complete hexamer of *Ng*SAT with four CoA and six serine bound in the active site was generated.

This merged structure was prepared for modelling using the protein preparation wizard (34, 35). Hydrogen atoms were added, and protonation states of ligands were assigned using Epik for a pH range of 7±2. The amino group of serine ligands were fixed to have a +1 charge, and the carboxylate terminal group of Gln264 residues was added. To remove steric clashes between merged substrates and the binding site residues, atoms on CoA and residues within 5 Å of the substrates were minimized using Prime energy minimization calculation. Then, hydrogen bonds were optimized, and protonation states of ionizable residues were assigned using PROPKA for pH 8.0. It should be noted that His162 was assigned a protonation state of HIS. An acetyl group was built onto each co-enzyme A molecule, followed by another round of Prime minimization for acetyl-CoA and residues within 5 Å of acetyl-CoA and serine.

In this new structure, there were two types of active site conformations: fully occupied active site (with both serine and acetyl-CoA) and partially occupied closed active site (with serine present but the acetyl-CoA site occluded by the closing C-terminal tail). The new complete structure with substrates bound was used to set up MD simulations. The MD system was built in Desmond (46, 47) by adding TIP3P explicit water molecules to an orthorhombic box. The box size was set by having a buffer region of 10 Å in each of the x, y, z directions. The box volume was minimized by rotating the enzyme molecule. Ions were added to neutralize the system, and NaCl salt was added at a concentration of 0.15 M. The force field used was OPLS_2005. MD simulation was conducted using Desmond (46, 47). The system was first minimized for 100 ps, followed by NPT ensemble MD calculations at a temperature of 310 K and a pressure of 1.01325 bar. Langevin thermostat and barostat were used. The MD simulation was conducted for 5 μs.

### Ligand library Preparation

Drug-like compounds with neutral charges were retrieved from the ZINC15 database (19). The 3D structures of these compounds, possessing drug-like properties, devoid of reactive groups, charge-neutral at pH 7, and available for purchase, were downloaded from the ZINC15 database (accessed on 28-08-2019). The initially downloaded library comprised approximately 7 million compounds. This compound set was subsequently filtered using Pan-assay interference compounds (PAINS) filters (48) and prepared for modelling in LigPrep (49), using the OPLS3 force field. Possible ionization states were generated for pH 7±1 using Epik, and potential tautomers were generated while retaining the specified chiralities in the compounds. The prepared library now consists of 10,479,301 compounds. For the large-compound library, 3D structures of compounds with a molecular weight greater than or equal to 500 Daltons, a logP value above 4, charge-neutral at pH 7, no reactive groups, and available for purchase, were downloaded from the ZINC15 database (accessed on 2021-06-21) (19). The initial library includes approximately 1 million compounds. The set of compounds was already filtered by PAINS filters upon download (48). The library was prepared using LigPrep with the OPLS4 force field. Possible states at pH 7±1 were generated using Epik; tautomers were generated while retaining chiralities within the 3D structures. The prepared library contains 1,294,740 compounds.

### Receptor grid generation

The substrate-bound active site model was used to generate the receptor grid for virtual screening. Firstly, the acetyl-CoA and serine substrates were removed. Two receptor grids were generated with slightly different positions for the center of the grid. For the first receptor grid, the center of the grid was defined as the centroid of residues 246, 231, 229, and 244 from chain A, and residues 246, 223, 205, 226, and 208 from chain C. For the second receptor grid, the center of the grid was defined as the centroid of residues 246, 231, and 244 from chain A, and residues 246 and 248 from chain C. The size of both grids was defined by allowing docking of ligands with a length <= 20 Å. For the first grid, rotation of the hydroxyl groups on Thr239 and Ser228 from chain C was allowed. For the second grid, rotation of the hydroxyl groups on Ser189 from chain A, and Ser228 and Thr239 from chain C was allowed. The receptor grids were generated using Glide (50-52)from the Schrödinger Suite (53).

During the 5 μs MD simulation for the full-length *Ng*SAT structure, the acetyl-CoA molecule bound in the interface between chains E and F remained stable; thus, this binding site was used for virtual screening of the large-compound library. A representative conformation (frame at 3.64 μs from the full trajectory) was extracted from the equilibrated time period of the MD simulation (3-5 μs) to be used as the receptor conformation for virtual screening. This conformation was prepared for receptor generation using the protein preparation wizard (34, 35). All water molecules were deleted, and hydrogen bond interactions as well as protonation states of ionizable residues were assigned using PROPKA for pH 8.0. Restrained minimization of the structure was then conducted to remove any steric clashes. A receptor grid was generated centering at the acetyl-CoA binding site between chains E and F. All other acetyl-CoA and serine ligands were deleted. The center of the grid was defined by the position of acetyl-CoA in the E/F interface. The size of the grid was determined by allowing docking of ligands with sizes similar to acetyl-CoA. Rotatable groups were allowed for side chains of Ser189 from chain F and Thr254 from chain E.

### Virtual screening

The prepared compound library was screened against the receptor grids generated above using the virtual screening workflow in Schrödinger Suite. Virtual screening (VS) was conducted in three docking stages with increasing precision, HTVS (high-throughput VS), SP (standard precision), and XP (extra precision). Top 10% ranked compounds from the HTVS stage were subject to SP docking, and the best 10% compounds from the SP stage were then subject to the final XP docking state. One pose was written out for each ligand, and the top ranked 1000 compounds were written out for evaluation.

### MM-GBSA calculation and MD validation

The docked poses of selected compounds, after examination of the virtual screening results, were further evaluated using MM-GBSA (molecular mechanics with generalized Born and surface area) calculations in Prime (40-42). MM-GBSA binding energies, representing approximate free energies of binding, were calculated as the difference between the energy of the ligand-enzyme complex and the sum of energies of the free ligand and free enzyme. A more negative value indicates a predicted stronger binding. MM-GBSA calculations were conducted using the OPLS4 force field and the VSGB solvation model. The binding poses were refined, and active site residues within 5 Å of the docked compound were minimized to optimize interactions.

After further examination of the MM-GBSA results, compounds that retained interactions with the enzyme and showed favorable predicted binding energies were selected for additional validation through short MD simulations (2 ns). The MD simulation system for each compound was constructed based on the MM-GBSA optimized binding pose, and explicit SPC water molecules were added to orthorhombic boxes. The box size was determined by establishing a buffer region of 10 Å in each of the x, y, and z directions. The box volume was minimized by rotating the enzyme-ligand complex. Na^+^ and Cl^-^ ions were incorporated to neutralize the system, and NaCl salt was added at a concentration of 0.15 M. The OPLS3e force field was employed to generate the MD system. MD simulations were performed using Desmond, with NPT ensembles at a temperature of 310 K and a pressure of 1.01325 bar. The system was relaxed using Desmond’s default protocol before the MD production run. Each MD validation calculation was run for 2 ns.

### Preparation of recombinant *Ng*SAT

*Ng*SAT was cloned and expressed using methods published previously (18). *Ng*SAT was cloned into expression vector pET28b-pstI and expressed in *E. coli* BL21 (DE3) with a N-terminal HexaHistag. *Ng*SAT was cultured in 1L of LB at 37°C and induced with Isopropyl β-d-1-thiogalactopyranoside IPTG (0.75 mM final concentration) once an 0.4-0.6 OD A_600_ was reached. Cultures were left to express at 37°C overnight (180 rpm) and pellets were stored at -80°C before purification. Thawed pellets were resuspended in lysis buffer (50 mM Tris pH 8.0, 200 mM NaCl, 20 mM Imidazole) with the addition of one cOmplete Mini EDTA-free protease inhibitor tablet (Roche). On ice, the resuspended pellet was sonicated using a ¼ inch probe, in one second bursts with one second intervals, for 1.5 minutes of sonication in total, followed by centrifugation for 20 minutes at 13,945 xg (10°C). For IMAC purification, 0.22 µM filtered supernatant was loaded onto a 5 ml HisTrap^TM^HP column (Cytiva) and eluted over a 50% imidazole gradient (50 mM Tris pH 8.0, 200 mM NaCl, 1 M imidazole) eluting at a final concentration of 400 mM imidazole. Eluted protein was concentrated at 3000 xg at 10°C using a 10K MWCO concentrator and loaded onto an analytical Enrich650 size exclusion column (Bio-Rad) pre-equilibrated in size exclusion buffer (50 mM Tris pH 8.0, 100 mM NaCl), where it eluted as a single peak with an elution volume of 12.9 ml consistent with a hexameric oligomer. Protein concentration was measured through measuring absorbance at A280nm (ε = 0.598) and diluted to an appropriate working stock concentration for *Ng*SAT activity assays and inhibitor screening. All assays were conducted in size exclusion buffer.

### *Ng*SAT activity assays and inhibitor screening

*Ng*SAT activity and inhibitor screening was measured in a 96-well plate format using a coupled DTNB assay monitoring the formation of CoA producing the colored product TNB detected at 412 nm (ɛ = 14,150 M^-1^ cm^-1^) (54). All assays were collected in 50 mM Tris pH 8.0, 100 mM NaCl at 25°C with 1 mM DTNB, 0.1 mM acetyl-CoA, 1 mM serine and 1.97 nM of purified *Ng*SAT, in a final reaction volume of 250 µl, unless otherwise stated. For inhibitor screening, substrate concentrations were kept at or below K_m_ (serine K_m_ = 1.21 mM, acetyl-CoA K_m_ = 0.15 mM (18)) to allow for detection of inhibitors with a competitive mode of inhibition. Reactions were measured for 20 minutes, and the initial linear velocity was determined from the linear portion of the reaction.

Inhibition assays were optimized through pre-incubation of the enzyme with inhibitors before initiating the reaction with a substrate mix of serine and acetyl-CoA. All compound stocks were prepared in 100% DMSO with 1-10% DMSO in the final reaction volume. All positive controls contained an appropriate amount of DMSO in the final reaction. All experimentally tested compounds were ordered with at least 90% purity from commercial vendors ChemBridge, MolPort and Asinex. Compound 2 was purchased from ChemBridge (Catalogue number 11296740, ZINC15 number ZINC11882369). *In vitro* screening of compounds at 100 µM was collected in the presence of 0.15 mM acetyl-CoA, 1 mM serine. IC_50_ dose response curves were collected in the presence of 0.1 mM acetyl-CoA and 0.5 mM serine through fitting the data to the non-linear regression dose-response curve – variable slope (Equation) using Prism (Graphpad, version 10.1.0).

## Equation 1. Inhibitor vs. response -Variable slope

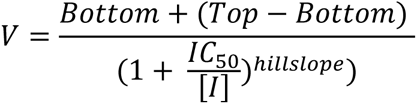

